# Identifying the neural representation of fast and slow states in force field adaptation via fMRI

**DOI:** 10.1101/582791

**Authors:** Andria J. Farrens, Fabrizio Sergi

**Author notes:** FS (corresponding author -), and AJF are with the Human Robotics Laboratory, Department of Biomedical Engineering, University of Delaware, Newark DE, 19713 USA.

## Abstract

Neurorehabilitation is centered on motor learning and control processes, however our understanding of how the brain learns to control movements is still limited. Motor adaptation is a rapid form of motor learning that is amenable to study in the laboratory setting. Behavioral studies of motor adaptation have coupled clever task design with computational modeling to study the control processes that underlie motor adaptation. These studies provide evidence of fast and slow learning states in the brain that combine to control neuromotor adaptation.

Currently, the neural representation of these states remains unclear, especially for adaptation to changes in task dynamics, commonly studied using force fields imposed by a robotic device. Our group has developed the MR-Softwrist, a robot capable of executing dynamic adaptation tasks during functional magnetic resonance imaging (fMRI) that can be used to localize these networks in the brain.

We simulated an fMRI experiment to determine if signal arising from a switching force field adaptation task can localize the neural representations of fast and slow learning states in the brain. Our results show that our task produces reliable behavioral estimates of fast and slow learning states, and distinctly measurable fMRI activations associated with each state under realistic levels of behavioral and measurement noise. Execution of this protocol with the MR-Softwrist will extend our knowledge of how the brain learns to control movement.

## I. Introduction

Neurorehabilitation is centered on the idea that retraining upper-extremity motor function following neural injury can be advanced by incorporating concepts of motor learning and motor control into treatment protocols. To effectively integrate these concepts into rehabilitation, motor learning and motor control strategies need to be better understood. Currently, how the brain learns to control movement remains an open question in the field of neuroscience. Motor adaptation, a rapid form of motor learning, can be used to study motor control in the laboratory setting.

Numerous behavioral studies have used motor adaptation as a tool to investigate neuromotor control processes [1]. Motor adaptation studies show that the brain appears to control movements by predicting the required task dynamics based on an internal model of the environment, and programs its motor output accordingly. When this prediction is incorrect, the resulting movement errors are used to update the internal model to generate updated predictive motor commands [1]. A plausible model that explains this adaptation control process is a multi-state system, with states that evolve at multiple timescales. Classically, this multi-state system has been modeled as a combination of one fast and one slow learning state, that are updated trial-to-trial by the magnitude of performance error [2], [3]. Recent work, however, suggests that a model with a single fast learning state, and multiple context dependent slow learning states better explain the behavioral data across multiple task schedules and adaptation modalities (force field, saccadic gain shift, and visuomotor transformation) [4].

While the behavioral effects of motor adaptation have been extensively studied, its neural correlates are much less understood. Previous works have tried to localize adaptation control processes (i.e. fast and slow states) in the brain by using a patient population with focal lesions or applying tDCS or TMS to map neural impairment to abnormal motor behavior [5]–[7]. While informative, these studies do not confirm or reject if the models that fit the behavioral results also explain the neural activity they stem from. Moreover, they provide only a broad mapping of brain region to motor function and can not assess the interactions between brain regions during adaptation. Instead, functional magnetic resonance imaging (fMRI) can be used to directly measure the neural correlates of motor adaptation as it occurs. fMRI is a whole brain imaging technique that measures metabolic activity in the brain with millimeter spatial resolution. Kim et. al recently used fMRI in visuomotor adaptation to provide evidence of distinct neural activations associated with slow and fast learning states [8]. While visuomotor adaptation exhibits similar behavioral traits as force field adaptation, it relies on different brain regions [9], [10]. Therefore, the neural representation of learning state networks responsible for force field adaptation remains an open question.

Using fMRI to determine the neural representation of force field adaptation learning states poses multiple challenges. The first is the development of a fMRI compatible system capable of executing a force field adaptation task in the MRI environment. Recently, our lab has developed the MR-Softwrist, a 3 degree of freedom wrist robot that has been validated as a tool for performing wrist pointing force field adaptation without causing fMRI image degradation [11]– [13]. The second challenges are inherent to fMRI, and include the low signal-to-noise ratio and the poor temporal resolution (≥1 sec) of the acquired signal, as well as estimation of neuronal activity from behavioral measurements that are prone to measurement error and subjects variability in motor execution [14].

In this paper, we verify whether simulated fMRI signal arising from exposure of subjects to a force field task, executable by the MR-Softwrist, has sufficient signal-to-noise ratio to identify the neural representations of fast and slow learning states estimated via a multi-state model. To test this, we designed a force field adaptation task capable of 1) producing behavioral measures that can be used to reliably estimate model parameters that define the evolution of internal states, and 2) producting distictly measurable fMRI activations associated with each internal state during task performance. We simulated behavior at four noise levels that modeled subjects behavioral variability and our system’s measurement error, and 3 realistic signal-to-noise levels reported in the literature [15]. Future use of our task with the MR-Softwrist will determine if the models that explain motor adaptation behavior also explain the neural activity it stems from.

## II. Methods

### A. Multi-state Model

Motor adaptation is the trial and error process of adjusting movement to new demands. The central nervous system (CNS) controls movement in a feed-forward manner based on its expectation, called an internal model, of the required movement dynamics. The CNS adapts to change in these dynamics through error feedback; movement (generated based on the internal model) that results in error is used to recalibrate the internal model and change subsequent motor output.

In order to simulate motor adaptation, we needed to chose a model that described both the behavioral and neural data previously reported in the adaptation literature. The model we chose is a multi-state model, proposed by Lee et. al [4]. Lee et. al. tested 10 candidate models to determine the model that could simultaneously account for the following motor adaptation phenomena commonly reported in the literature: savings, spontaneous recovery, anterograde interference, and dual adaptation in both blocked and random schedules. They identified a multi-state model with 1-fast and n-slow states acting in parallel as the simplest model capable of accounting for all behavioral phenomena. The fast state is a single state, while the slow state has multiple context-dependent inner states. The model for this system is as follows:

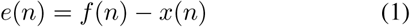

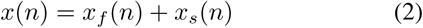

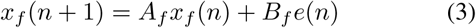

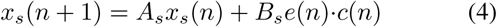

In this model, performance error *e*(*n*) is modeled as the difference between the applied force field *f* (*n*) and the subjects motor output (controlled by the CNS internal model) *x*(*n*) (eq. 1). Here, *x*(*n*) represents the internal model on trial (n), that is comprised of two states (eq. 2), a fast learning state *x*_*f*_ and a slow learning state *x*_*s*_, organized in a parallel architecture. The fast learning state is updated on every trial by performance error *e*(*n*) (eq. 3). The slow learning state has *N*_*condition*_ inner states, equal to the number of dynamic conditions included in the adaptation task (eq. 4). Each inner slow state is updated by performance error *e*(*n*) only for its corresponding task condition, and is engaged non-linearly by a contextual cue *c*(*n*). For example, for the first task condition, (c)=[1, 0,…0], and for the second task condition, (c)=[0,1,… 0], and so on. In this way, the slow states are updated by error only during their respective task conditions. *A*_*f*_, and *A*_*s*_ are constants that represent the retention factor of each state (i.e. how much the previous state contributes to the current state), while *B*_*f*_ and *B*_*s*_ define the sensitivity of each state to error. For these parameters, *A*_*f*_ < *A*_*s*_ and *B*_*f*_ > *B*_*s*_ as the slow state retains more from trial to trial, and is less sensitive to error than the fast state.

### B. Task Design

Recent work in visuomotor adaptation used a switching task performed during fMRI to provide evidence of distinct fast and slow learning state networks in the brain [8]. This work was exploritory in nature, and did not use a pre-defined model of expected neural activations associated with adaptation in their analysis. Moreover, evidence from the literature suggests that visuomotor and force field adaptation rely on seperate neural processes-a finding that seems apparent when considering the different modalities of proprioception involved in performing each task [9], [10]. Consequently, the aim of this paper is to design a force field adaptation task capable of 1) producing behavioral measures that can be used to reliably estimate model parameters (*A*_*f*_, *A*_*s*_, *B*_*f*_, *B*_*s*_) that define the evolution of internal states, and 2) that produces distictly measurable fMRI activations associated with each internal state during task performance.

While simulated in this work, we plan to execute the designed task using the MR-Softwrist. As such, the premise of our task is that subjects will control the handle of the MR-SoftWrist to move a cursor displayed on a monitor. Radial ulnar deviation (RUD) of the wrist will move the cursor vertically while flexion-extension (FE) will move the cursor horizontally (rotation about the radio-ulnar axis will be prevented by a support). Subjects will be cued to make alternating FE wrist rotations to move the cursor along a straight path to one of two targets, placed at (±12, 0) degrees in FE, RUD respectively. To induce force field adaptaiton, we will use a standard curl-force field, in which the robot applies a velocity-dependent torque, *τ* = *B* ·*θ*, where *τ* = [*τ*_*FE*_, *τ*_*RUD*_], is proportional to the measured velocity 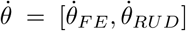 scaled by magnitude *B* = [0, 1; −1, 0] mN·m·s/deg [1], [13].

According to the multi-state model, the engagement of slow learning states is dependent on the number of force field conditions included in the task (eq. 4). For our experiment, we chose to use four force field conditions: Conditions A and B are clockwise and counterclock curl force fields with magnitudes ± 1·B respectively, while conditions C and D are curl force fields with magnitudes ±2 ·B respectively. To provide a contextual cue (*c*(*n*)) for subjects to distinguish between condition type, change in task condition will be cued by a change in target color.

Our task is divided into two phases as shown in Fig. 1. In Part 1, subjects practice conditions A and B in a blocked schedule, that consists of three consecutive blocks of 60 trials of condition A, 60 trials of condition B, and 48 trials of condition A. Part 1 is used to produce behavioral measurements that can be used to reliably estimate model parameters. In Part 2, subjects perform an alternating schedule of conditions C and D, that was designed to elicit distinctly measureable fMRI activations.

**Fig 1.**
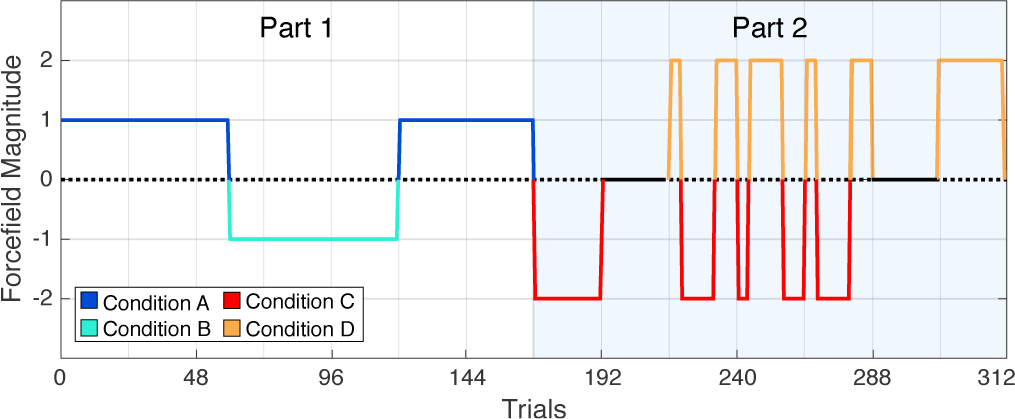
Task Design. Part 1 is used to estimate model parameters, Part 2 is designed to maximally dissociate neural activations associated with inner slow learning states

### C. Simulated Experiment

To determine if our designed task is feasible for use in an fMRI study aimed at dissociating neural activations associated with fast and slow learning states, we simulated our experiment (Fig. 2). We chose parameters *A*_*f*_, *A*_*s*_, *B*_*f*_, *B*_*s*_ from the literature and used the multi-state model to simulate 1) a subjects behavioral output and 2) the evolution of the corresponding fast and slow learning states. We then added experimental noise to each measure, and used the noisy data to re-estimate state-specific regressors. In parallel, the output of the model was used to simulate fMRI data at three noise levels taken from the literature. We thus performed analysis of the simulated fMRI data based on estimated learning states taken from our simulated behavioral measures to determine if the resulting behavioral measures and corresponding fMRI signals could be used to reliably identify state-specific neural representations.

**Fig 2.**
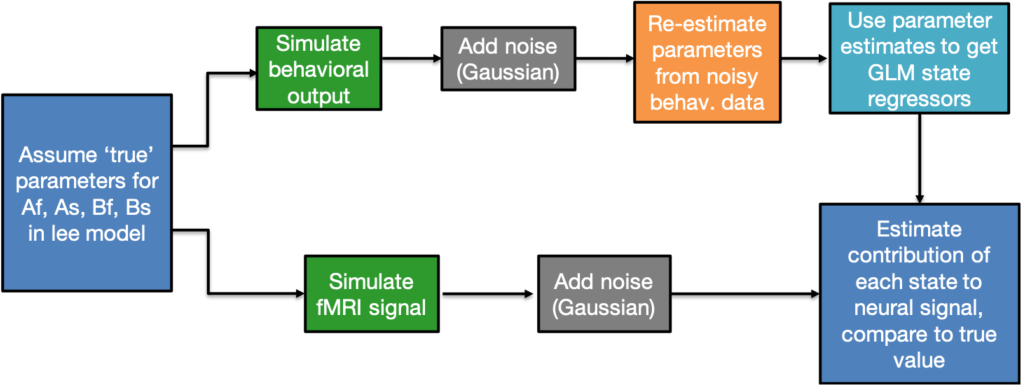
Flow chart of simulated experimental design

#### 1) Behavioral Simulation

To create our simulated data sets, we chose model parameters *A*_*f*_ = 0.825, *A*_*s*_ = 0.9901, *B*_*f*_ = 0.3096, *B*_*s*_ = 0.2147 to generate *true* behavior assuming zero noise [4]. From this *true* data set we created two seperate behavioral datasets based on two different behavioral measurements, and added task-appropriate noise.

In force field adaptation, the internal model, *x*(*n*), can be estimated from behavioral measurements in two ways: continuously, via measured trajectory error (TE), or intermittently over the course of training via an adaptation index (AI). For TE, the internal model is estimated using eq. 1, through knowledge of the applied force field *f* (*n*) and measured TE as an approximation for *e*(*n*). TE is typically taken as the internal angle between the cursor at the peak velocity during the initial ballistic movement and the starting target relative to the end target (Fig. 3). In this way, TE is measured with limited interference from online error correction processes and most directly reflects the feed-forward commands based on the current state of the internal model. TE is normalized by the first TE measured in response to perturbation, when the internal model *x*(*n*)= 0, such that all TE measures are a percentage of initial TE. As TE is measureable on every trial, the simulated TE behavioral set was created for every trial.

**Fig 3.**
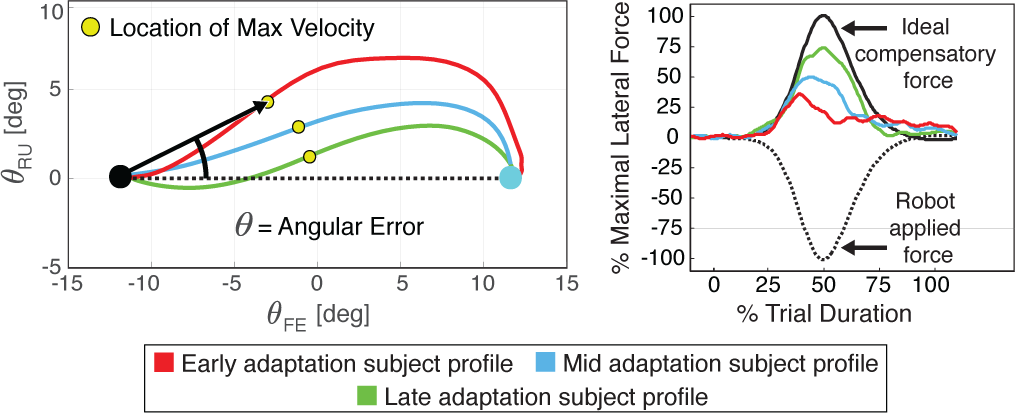
Behavioral measurements used for internal model estimation

**Fig 4.**
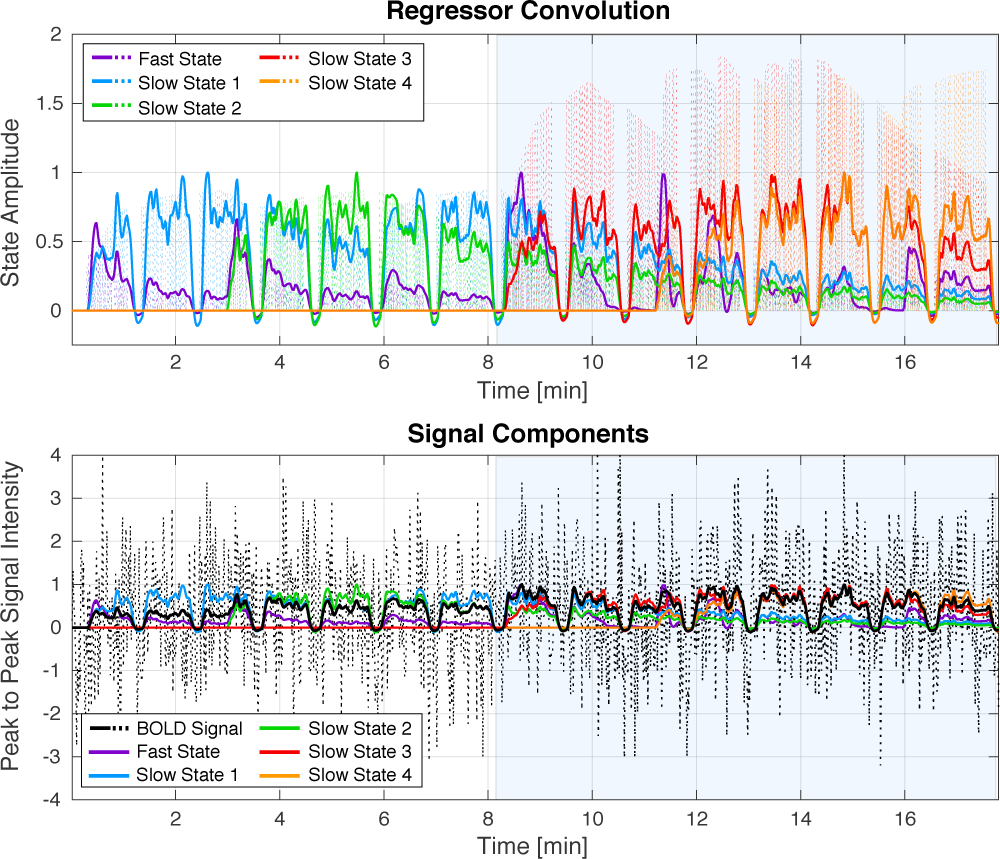
Top: Dashed lines show stimulus functions of each learning state, solid lines depict normalized regressor signals generated after convolution (s(t)*h(t)). Bottom: Contribution of each learning state to overal signal fluctuation (1%) with added noise corresponding to CNR = 1.2

AI is measured during error clamp trials, in which lateral trajectory errors are ‘clamped’ to 0 such that *x*(*n*)= *f* (*n*). In this condition, *f* (*n*) is the lateral force profile produced by the subject that is measured by the robot. This lateral force profile reflects the output of the subjects internal model, and is a method of sampling the internal model that is free from online error corrections as performance error is clamped to 0, although this is dependent on the robot’s ability to apprioriately ‘clamp’ errors. AI is determined by regression between the measured force profile and the ideal force profile to produce a normalized estimate of *x*(*n*) (Fig. 3). Because the perturbing force field is removed during error clamp trials, they can only be measured occasionally so as to not disrupt ongoing adaptation. Consequently, the AI behavioral set was sampled with a 1/8 frequency in a psuedorandom manner that ensured samples were atleast 4 trials apart [16].

Both behavioral measurement methods are subject to two different forms of variability. The first is subject’s inherent motor variability in task execution (process noise). The second arises from the resolution and accuracy of our measurement system that imparts a certain amount of measurement error to all behavioral measures. To incorporate an appropriate estimate both components of variability in our simulations, we had 12 subjects perform 144 trials of wrist pointing with the MR-Softwrist acting in a transparent no force condition, and sampled their TE and AI using the same psuedorandom schedule described above. We took the standard deviation of each subject’s behavioral measures across all sampled trials, and averaged the standard deviation across subjects to estimate the expected variability in each measurement type. For TE, this estimate was *σ*_*TE*_ = 0.12 and for AI it was *σ*_*AI*_ = 0.20 for each normalized measure. The higher variability in our AI measure is likely due to the compliance of the MR-Softwrist. The mean of both measures was zero, suggesting no directional bias in our system. Therefore, we assumed noise would follow a gaussian distribution with a mean of zero and the respective computed standard deviations. To test the robustness of our estimation methods to noise, we conducted simulations at the following noise levels [*σ*_*TE*_, *σ*_*TE*_ + 0.1, *σ*_*TE*_ + 0.2, *σ*_*TE*_ + 0.3] for the TE behavioral dataset, and [*σ*_*AI*_ − 0.1, *σ*_*AI*_, *σ*_*AI*_ + 0.1, *σ*_*AI*_ + 0.2] for AI. The reduced noise condition in AI was tested to see how improvements in our AI measure could affect parameter estimation.

To determine the feasibility of estimating the *true* model parameters given the available behavioral datasets, we fit the multi-state model to the behavioral data in Part 1 of our task. We created 100 behavioral data sets for each combination of noise condition and behavioral measurement type. For each data set the best-fit model parameters were estimated, to enable creation of confidence intervals for our parameter estimates in each simulated experimental condition. To find the parameter estimates that best fit the simulated data, we used a Markov Chain Monte Carlo (MCMC) method that ran for 150,000 iterations to search the model parameter space [17]. Prior distributions were defined using the mean and standard deviation of the model parameters reported in the literature (*Af* = 𝒩(0.7817, 0.1643); *As* = 𝒩 (0.9947, 0.0041); *Bf* = 𝒩(0.1515, 0.1433); *Bs* = 𝒩(0.1019, 0.1061)) [4]. *Af* was bounded betweem 0.5 and 1, *Bf* was bounded between 0 and 0.5. All slow state parameters were bounded between 0 and 1. For each iteration (i.e. parameter combination explored), the log-likelihood of the model fit to the behavioral data set was evaluated according to eq. (5).

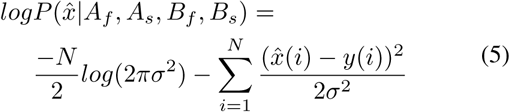

In this equation, 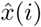 is the behavioral estimate of *x*(*i*) on trial *i* (using TE or AI), *N* is the total number of trials sampled, and *y*(*i*) is the model prediction of *x*(*i*) on the *i*^*th*^ trial. *σ*^2^ is the variance of the model output that includes the variance of behavioral output and state noise [4]. As an estimate of *σ*^2^, we simulated 1000 behavioral data sets of the corresponding measurement type and noise level being evaluated, and calculated the average sample variance as: 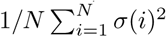, where *σ*(*i*)^2^ was the sample variance of the data on the *i*_*th*_ trial across 1000 iterations [4]. For each behavioral dataset, the parameter estimate with the maximum log-likelihood over all MCMC iterations was selected. The median parameter estimates arising from each condition (noise and measurement type) was then used in the multi-state model to create estimates of the trial-by-trial evolution of the fast 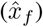 and slow states 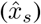 over the entire task.

#### 2) fRMI Simulation

The blood oxygenation level-dependent (BOLD) signal acquired during fMRI stems from changes in deoxyhemoglobin concentration in the brain, that occur due to stimulus related neuronal activity. BOLD signal is commonly modeled by convolution between a task-related stimulus function *s*(*t*) and the hemodynamic response function (HRF) *h*(*t*) such that *f* (*t*)= *s*(*t*) * *h*(*t*) [18], [19]. To create our simulated fMRI signal, *f* (*t*), we used SPMs standard HRF function with a TR = 1 [20].

To create our stimulus functions, the *true* states (*x*_*f*_ and *x*_*s*_) and the estimated states (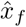 and 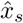) were converted into the time domain to create the functions *x*_*f*_ (*t*) and 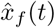, and 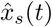. This was achieved by creating a pulse function that assigned the state value at a given trial to an appropriate onset time, estimated using realistic trial durations (*t*_*trial*_) and inter-trial intervals (*t*_*IT* *I*_): *t*_*trial*_ = 𝒩(0.5, 0.15) s; *t*_*trial*_ *>* 0.25 s, *t*_*IT* *I*_ = (2, 0.75) s; 0.5 s *< t*_*IT* *I*_ *<* 3 s. We included 15 sec. rest blocks between each block of 24 trials and between Part 1 and 2 of the task, both for subject comfort and to allow for relaxation of HRF.

The stimulus functions *x*_*f*_ (*t*) and *x*_*s*_(*t*) were convolved with the HRF and sampled with a TR of 1 s to create simulated fMRI datasets of measured BOLD signal. The simulated fMRI signal was generated to have a 1% peak to peak signal change [18], [19]. To model physiological and thermal noise present in real fMRI data, we added 3 levels of guassian noise (low, medium, high) to our simulated signal that correspond to temporal contrast to noise ratios of (1.2, 0.8, 0.5) [15].

Our estimated states, 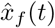 and 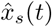, were used to create a General Linear Model (GLM) for regression onto our simulated fMRI data. Each estimated state was convolved with the HRF, and included in a GLM. We defined our first GLM (GLM 1) as:

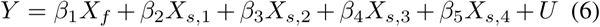

To test our ability to detect each state independently using GLM 1, we simulated five signals (1-5) with signal change originating from one of each of the five learning states. We created two signals with signal change arising from multiple neural stimuli, modeled as the summation of all slow states (signal 6), and all states (signal 7) [19]. GLM 1 was used to test our ability to detect individual states contributions to BOLD signal arising from the combination of multiple states. An additional GLM (GLM 2), defined as

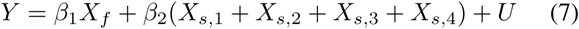

was used to test our ability to dissociate between signal arising from the fast state or the conglomerate of all slow states in signals 6 and 7. This was motivated by the literature, which suggests the neural representation of slow learning states may not be spatially distinct [8]. Under this assumption, regions responsible for slow state learning would generate signal reflective of the combination of all slow states rather than each individual slow state.

## III. Results

### A. Parameter Estimates

Box plots of the maximum log-likelihood parameter estimates found via MCMC at each noise level are reported in Fig. 5. For all TE noise levels, estimation of *A*_*f*_ had ≤ 2% error compared to the *true* value. For AI estimates of *A*_*f*_, percent error ranged from 0.7-10%. Percent error of median parameter estimates for *A*_*s*_ was less than 1% for all conditions. Both behavioral measurement types had larger percent error in the sensitivity parameter (*B*) estimations. For TE estimates of *B*_*f*_, percent error ranged from 2.5-17%, and for AI estimation error ranged from 9-32%. For TE estimates of *B*_*s*_, percent error ranged from 3-11%, and for AI estimation error ranged from 30-52%. Expectedly, as noise level increased the percent error of parameter estimates increased, as did the variability in parameter estimation, visualizable in Fig. 5. Across all noise conditions, the AI measure had larger variability in parameter estimation compared to the TE measure, and a larger estimation bias in error sensitivity parameters. Misestimations in parameter estimates led to larger error in fast and slow state estimations for the AI condition, as shown in Fig. 6.

**Fig 5.**
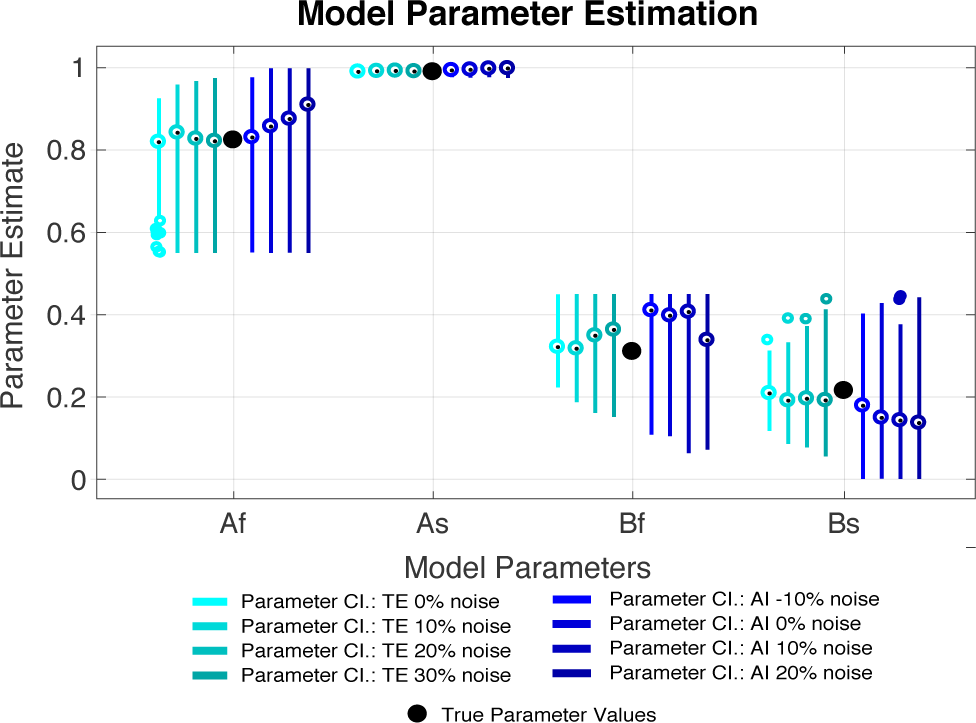
Boxplots of estimated model parameters for each behavioral dataset

**Fig 6.**
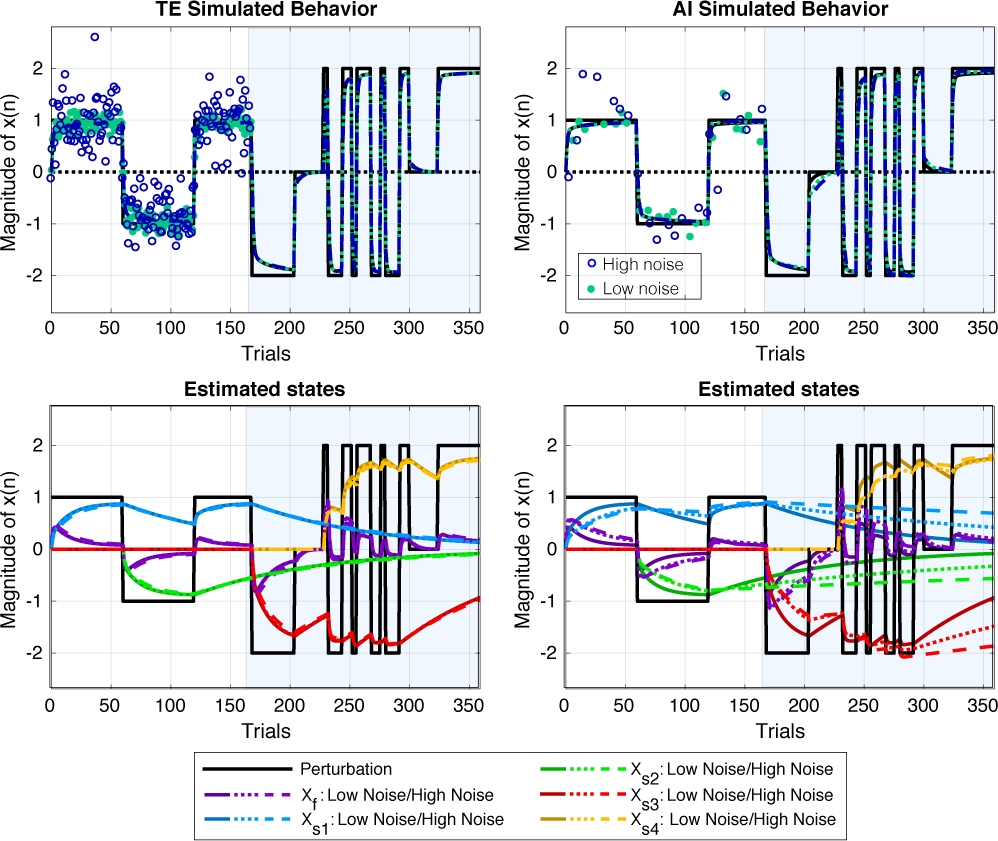
The behavioral estimation results for the lowest and highest noise conditions. The top figure shows the data used to estimate model parameters, and the bottom figure shows the *true* evolution of learning states in solid lines, and the estimated evolution of learning states in low and high noise conditions (low=short dashes, high= long dashes)

### B. fMRI Simulation Results

The sensitivity of our GLMs in identifying the correct learning states in each fMRI signal tested was quantified in terms of true positive (TPR) and false positive (FPR) rates, displayed in Fig. 7. At the CNR = 1.2 noise level, for signals 1-5, GLM 1 identified signal associated with each individual state regressor with a *>*98% TPR in all behavioral noise conditions. At the CNR = 0.8 noise level, GLM 1 had a >90% TPR in identifying individual learning states, except for behavioral conditions σ_*AI*_ + 0.1 and σ_*AI*_ + 0.2. At the CNR = 0.5 noise level, GLM 1 had a >50% TPR in identifying the fast learning state in signal 1, and >70% TPR for slow states (signals 2-5), excluding the σ_*AI*_ + 0.1 and σ_*AI*_ + 0.2 conditions.

**Fig 7.**
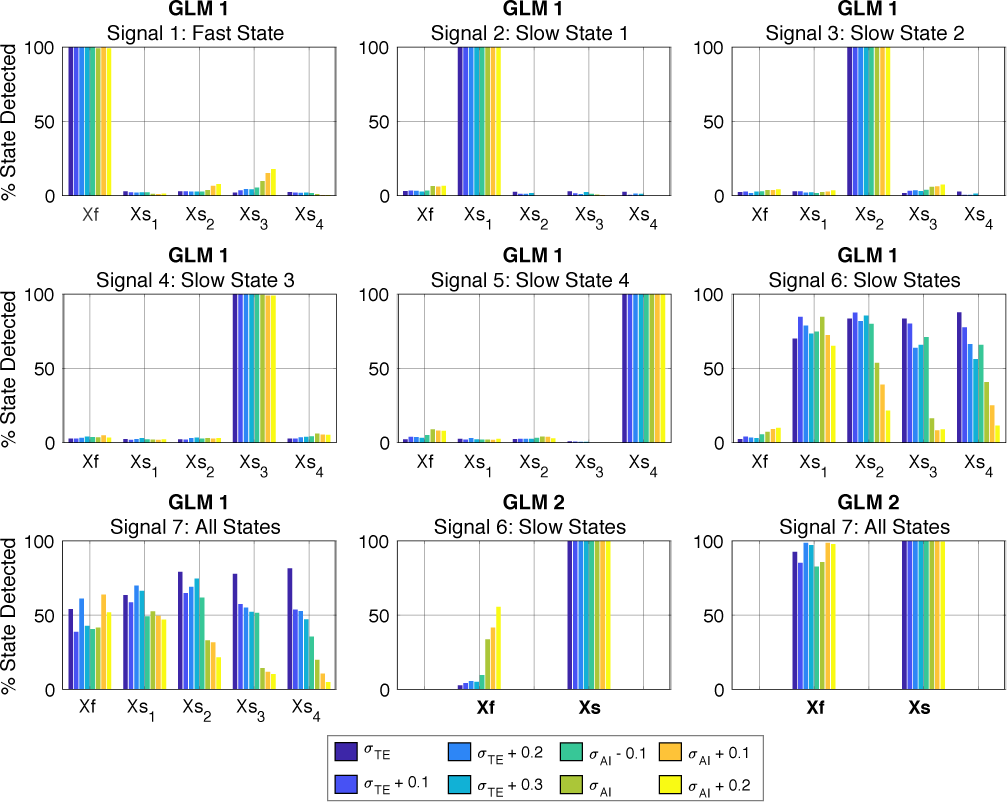
GLM parameter estimates in the CNR=1.2 noise condition

GLM 1 was less accurate in identifying individual state activations in fMRI BOLD signals that arose from combinations of learning state activations. For signal 6, at CNR = 1.2, GLM 1 identified each state with >50% TPR, except in behavioral conditions *σ*_*AI*_ +0.1, and *σ*_*AI*_ +0.2. For signal 7, at CNR = 1.2, GLM 1 identified each state with *>*50% TPR only in the lower noise TE conditions (Fig. 7). At higher noise levels (CNR = 0.5 or 0.8), in signals 6 and 7, GLM 1 was less than 50% accurate in identifying individual learning state contributions across all behavioral noise conditions.

The prevalent misidentification of the slow states in the AI conditions was largely due to the mis-estimation of the error sensitivity parameters, as the lower *B*_*s*_ estimates produced regressors that were more highly correlated with each other, especially for regressors *X*_*s*,3_ and *X*_*s*,4_.

GLM 2 was more accurate in dissociating fast and slow learning states in combination signals 6 and 7. For signal 6, at CNR = 1.2, GLM 2 identified the combined slow learning states activation with 100% TPR for all behavioral conditions. However, an elevated FRP (30-45%) in fast state detection was seen in behavioral conditions σ_*AI*_, σ_*AI*_ + 0.1, and σ_*AI*_ + 0.2. At higher noise levels, identification of the slow state remained >90% TPR for all behavioral conditions, while the fast state FPR decreased. For signal 7, at CNR = 1.2, GLM 2 identified the fast state with >80% TPR, and the combined slow learning states with 100% TPR for all behavioral conditions. At CNR = 0.8, this dropped to >50% and >90%, and to *<*50% and >50% for CNR = 0.5.

## IV. Discussions and Conclusion

The results of our simulation show that our switching force field task is capable of detecting fMRI signal arising from fast and slow learning states, determined via fitting a multi-state model to behavioral measurements of trajectory error or adaptation index under realistic behavioral and fMRI noise conditions. Using GLM 1, our task is powered to identify regions associated with each individual learning state. While GLM 1 was underpowered to dissociate state specific activations in signal arising from combinations of learning state activity, GLM 2 was able to detect activations associated differentially with the fast state or combined slow state regressor, that reflects the expected neural activations arising from brain regions responsible for slow state learning. Our simulaton further indicates that TE is a better method for model parameter estimation, however this result may underestimate the effects of online-error correction processes annal d joint stifening that could reduce the efficacy of this measure. Our next steps will be to re-evaluate our test design under longer experimental conditions to increase state detection power for fMRI signal arising from combinations of states, and to apply this methodology in-vivo with the MR-SoftWrist.

## V. Acknowledgments

We acknowledge support from the American Heart Association Scientist Development Grant no.17SDG3369000.

